# Multiple roles for TFG ring complexes in neuronal cargo trafficking

**DOI:** 10.1101/2024.11.05.621662

**Authors:** Ziheng Zhang, Molly M. Lettman, Amber L. Schuh, Basudeb Bhattacharyya, Peter Randolph, Trisha Nandakumar, Ishan Kulkarni, Alexander Roach, James R. Alvin, Diane Gengler, Scott M. Stagg, James L. Keck, Anjon Audhya

## Abstract

Pathological variants in Trk-fused gene (TFG) have been implicated in a variety of neurodegenerative conditions. In particular, mutations within its amino-terminal PB1 domain have been suggested to cause hereditary spastic paraplegia (HSP), resulting in progressive lower limb spasticity and weakness. The structural basis for this effect is unknown. Here, we combine X-ray crystallography and cryo-electron microscopy to determine a structural model of TFG, demonstrating the mechanism by which it forms octameric ring complexes. A network of electrostatic and hydrophobic interactions defines the interface between protomers. Moreover, we show that mutations identified previously in HSP patients disrupt this interface, destabilizing octamers, which ultimately leads to axonopathy. Surprisingly, the impacts of these variants are not equivalent in vivo, highlighting the existence of multiple, distinct mechanisms by which TFG mutations contribute to neurodegenerative disease.

## Introduction

The corticospinal tract (CST) is one of the major neural pathways for directing voluntary movement in primates, transmitting information from the cerebral cortex through the spinal cord to enable fine control of muscles^1,2^. Of particular importance are glutamatergic pyramidal neurons that form the CST, descending from layer V of the cortex and sometimes reaching up to a meter in length^3^. Pathological mutations or external injury that damage these neurons often lead to deficits in motor control^4^, but the mechanisms that underlie neurodegeneration following such insults remain poorly understood. By studying monogenetic neurological diseases however, specific pathological impacts to neurons can be revealed, yielding crucial insights into their normal function. Hereditary spastic paraplegias (HSPs) are typically single-gene disorders and represent one of the most common inherited neuropathies, collectively affecting hundreds of thousands of patients worldwide^5,6^. Importantly, the array of genomic loci implicated in HSP has successfully illuminated key roles for membrane transport, lipid metabolism, and myelination in neuronal maintenance^7^.

Mutations in Trk-fused gene (TFG) have been suggested to predispose neurons to degenerate, both in the central and peripheral nervous systems, resulting in a wide range of disease states, including HSPs, amyotrophic lateral sclerosis (ALS), Charcot-Marie-Tooth disease type 2 (CMT2), hereditary motor and sensory neuropathy (HMSN), and parkinsonism^8–13^. In general, pathological variants within the amino-terminal PB1 (Phox and Bem1) and coiled coil domains are associated with early onset forms of HSP, while those in the carboxyl-terminal disordered region result in adult-onset peripheral neuropathies^14^. Several of these mutations have been suggested to impact the ability of TFG to reversibly undergo liquid-liquid phase separation (LLPS), leading to aggregate formation in some cases, while others impede condensate formation^9,10,15–17^. However, the mechanisms that direct TFG phase transitions remain largely unknown, at least in part due to a dearth of high resolution structural information needed to guide atomistic or coarse grained modeling approaches.

Based on negative staining electron microscopy, TFG was shown previously to form 11-nm octameric cup-like structures that exhibit a propensity to self-associate in solution, depending on ionic strength, creating particles that are approximately 200-300 nm in diameter, similar in size to TFG condensates found in cells^18^. A recessive mutation in the TFG coiled coil domain (p.R106C) that underlies HSP disrupts the conformation of octamers and impairs the ability of TFG to undergo phase separation in numerous cell types, including cortical neurons^19^. Similarly, a mutation in the amino-terminal TFG PB1 domain (p.R22W), also associated with HSP, was shown to perturb the assembly of TFG cup-like structures in vitro, although its impact on TFG distribution in cells remains unknown^19,20^. By contrast, dominant-acting pathological variants identified in the TFG carboxyl-terminus (p.G269V or p.P285L) cause TFG to form amyloid-like fibrils in vitro, which accumulate with aggregated TDP43 in the frontotemporal lobe of HMSN patients^9,10,17,21^. Thus, depending on the functional domain of TFG impacted, differing phenotypic consequences can result, highlighting the existence of multiple distinct mechanisms by which TFG mutations lead to neurodegenerative disease.

Based on both live cell imaging and fixed cell analysis, TFG localizes mainly to the early secretory pathway, between the endoplasmic reticulum (ER) and ER-Golgi intermediate compartments (ERGIC), although more recent studies have highlighted an additional pool of TFG associated with endosomes, particularly in axons and dendrites^16,18,22–24^. While its accumulation at the ER-ERGIC interface depends upon an interaction between the TFG carboxyl-terminus and components of the COPII coat^24^, how TFG targets to endosomes remains unknown. Moreover, the relative contributions of the TFG PB1 domain to promoting octamer assembly as compared to its role in directing TFG to sites of action remain unclear^22^. Here, we report the 1.91 Å resolution crystal structure of the TFG PB1 domain, demonstrating that its acidic OPCA (OPR/PC/AID) motif forms intermolecular salt bridges with a series of basic residues near its amino-terminus, consistent with a Type I/II fold. Additionally, key hydrophobic interactions further stabilize the protomer interface. By docking the atomic structure into a map resolved using cryo-EM, we establish the mechanism by which TFG forms octameric ring complexes and reveal how various pathogenic mutations in TFG disrupt its cellular function. Surprisingly, mutations in the PB1 and coiled coil domains of TFG result in distinct perturbations to its stability and localization specifically in neurons, despite both resulting in motor phenotypes associated with HSP. These data strongly suggest additional roles for the TFG PB1 domain beyond promoting octameric ring assembly in the central nervous system (CNS).

## Results

### Structural determination of TFG octameric ring complexes

Using small angle X-ray scattering, we previously showed that the TFG amino-terminus (residues 1-96) forms multimers in solution, precluding high resolution structural determination^19^. To circumvent this problem, we incorporated two point mutations (D60A and D62R) into the OPCA motif, analogous to those made to determine the structure of the p62/sequestosome PB1 domain^25^, which facilitated crystallization. We subsequently used molecular replacement to determine the TFG PB1 domain structure to 1.91 Å resolution (**Figure 1A and Table 1**).

**Figure 1.**
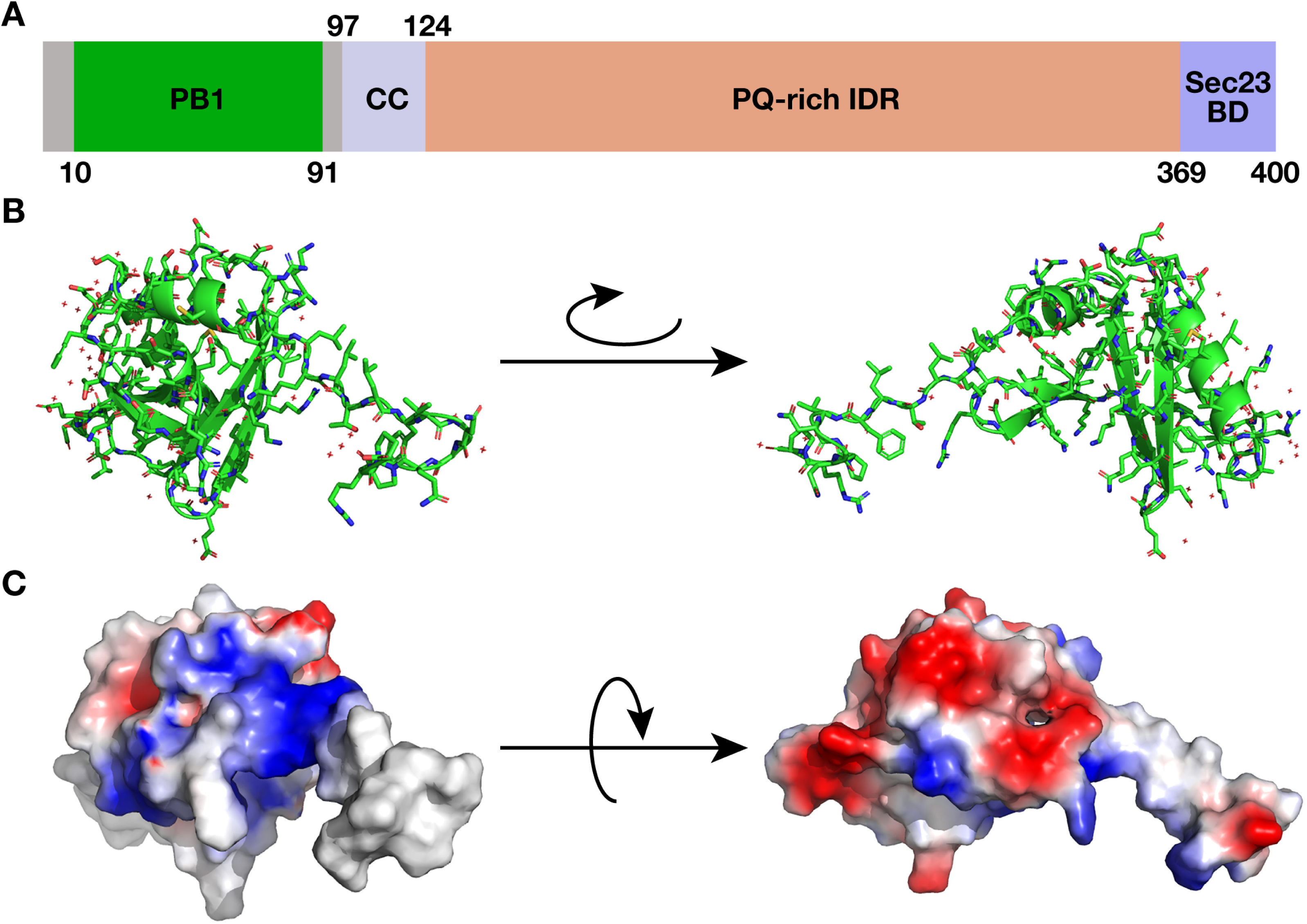
The TFG PB1 domain structure. (A) Illustration highlighting the domains of the human TFG protein (PB1, Phox and Bem1; CC, coiled coil; IDR, intrinsically disordered region; BD, binding domain). (B) Cartoon representation of the TFG PB1 domain structure (oxygen atoms, red; nitrogen atoms, blue). (C) Simulated surface protein contact potential of the TFG PB1 domain generated by PyMol (negative charge, red; positive charge, blue).

**Table 1.** X-ray Data Collection and Structure Determination Statistics.

|  |  |
| --- | --- |
| Data Collection |  |
| Wavelength, Å | 0.97872 |
| Resolution Range (high resolution bin), Å | 50-1.91 (1.94-1.91) |
| Space Group | C2 |
| Unit Cell (a, b, c (Å)) | 107.02, 32.49, 29.25 |
| ($\alpha$, $\beta$, $\gamma$ (°)) | 90.0, 104.8, 90.0 |
| Completeness, % | 99.4 (97.5) |
| Total/Unique Reflections | 7689/2480 |
| Redundancy | 7.2 (6.1) |
| $\langle I/\sigma I \rangle$ | 26.9 (2.2) |
| $R_{\text{sym}}^{\dagger}$ | 0.071 (0.536) |
| Refinement |  |
| Resolution, Å | 50-1.91 |
| $R_{\text{work}}/R_{\text{free}},^{\ddagger}$ | 0.2107/0.2534 |
| Rms deviations |  |
| Bonds, Å | 0.0096 |
| Angles, ° | 1.3514 |
| Ramachandran statistics, % |  |
| Most favored | 91.1 |
| Allowed | 8.9 |
| Generously allowed | 0 |
| Disallowed | 0 |
| # atoms |  |
| protein | 713 |
| solvent | 58 |
| $\langle B \text{ factor} \rangle, \text{Å}^2$ | |
| protein | 35.1 |
| solvent | 38.2 |

$\langle I \rangle$ is the mean intensity for multiply recorded reflections.
$^{\dagger} R_{\text{sym}} = \sum \sum_j |I_j - \langle I \rangle| / \sum I_j$ , where $I_j$ is the intensity measurement for reflection $j$ and $\langle I \rangle$ is the mean intensity for multiply recorded reflections.
$^{\ddagger} R_{\text{work}}/R_{\text{free}} = \sum ||F_{\text{obs}}| - |F_{\text{calc}}|| / |F_{\text{obs}}|$ , where the working and free R factors are calculated by using the working and free reflection sets, respectively. The free R reflections (5% of the total) were held aside throughout refinement.

Similar to other Type I/II PB1 domains, the TFG PB1 exhibits a ubiquitin like β-grasp fold composed of two α-helices and six β-strands (**Figure 1B**). Crystals included a single monomer in the asymmetric unit with well-resolved electron density for residues 10-96. While residues 10-82 fold as a globular domain, residues 83-96 are swapped with a crystallographic symmetry mate, contributing a β-strand within the core of the adjacent protein (**Figure 1B**). Based on a prediction of the domain structure using AlphaFold, β-strand swapping was not expected. Notably, similar artifactual β-strand swapping in structures of isolated protein domains has been observed in other proteins^26^.

The TFG PB1 domain harbors two dominant faces that are oppositely charged (**Figure 1C**), highlighting a potential mechanism of intermolecular dimerization. Based on prior negative staining EM-based studies, the coiled coil motif found immediately downstream of the PB1 domain was suggested to organize TFG protomers into octameric cup-like structures^18^. To enhance our understanding of how these complexes form, we used cryo-EM to generate a density map of the amino-terminal half of TFG (residues 1-193). Based on 2D classification of cryogenically prepared particles, we found that this region of TFG forms an 8-membered ring (**Figure S1A**). Class averages showed densities for secondary structures, indicating that the TFG particles can align to high resolution. However, there were no indications of side views in the 2D averages, which prevented us from performing a 3D reconstruction. We explored numerous specimen preparation parameters to try and capture side views, including continuous amorphous carbon, graphene, graphene oxide, carbon holey grids of multiple types, gold holey grids, nanoliter dispensing with rapid freezing (SPOTITON), and a variety of glow discharging parameters. None of these produced satisfactory side views, but the SPOTITON sample produced the best 2D class averages. Thus, we collected a tilted dataset on the SPOTITON grid, but the resulting class averages appeared blurry, indicating that the ice was becoming too thick with the tilted data for high precision particle alignment (**Figure S1A**). The best particles from the untilted and tilted datasets were combined, and a 3D single particle refinement was performed resulting in a 6.8 Å map.

Given the lack of side views, the resolution is highly anisotropic (**Figure S1B-S1D**), but we were able to unambiguously dock a modified PB1 domain structure where residues 83-96 are folded with a monomeric structure (**Figure 2A**). The resulting model demonstrated a coordinated series of salt bridges and hydrophobic interactions, positioning K14 on β strand B1 of one subunit adjacent to the OPCA motif of another in a front to back manner (**Figure 2A and 2B**), similar to PB1 domains found in p62 and atypical PKC^27^. Both the EM maps and modeling were inconsistent with the β-strand swap observed in the crystal structure, suggesting that the swap does not contribute to interactions between TFG PB1 domains. Strikingly, several pathological mutations associated with HSP were identified at this interface, suggesting a common mechanism by which they disrupt TFG function and lead to neurodegeneration (**Figure 2C**).

**Figure 2.**
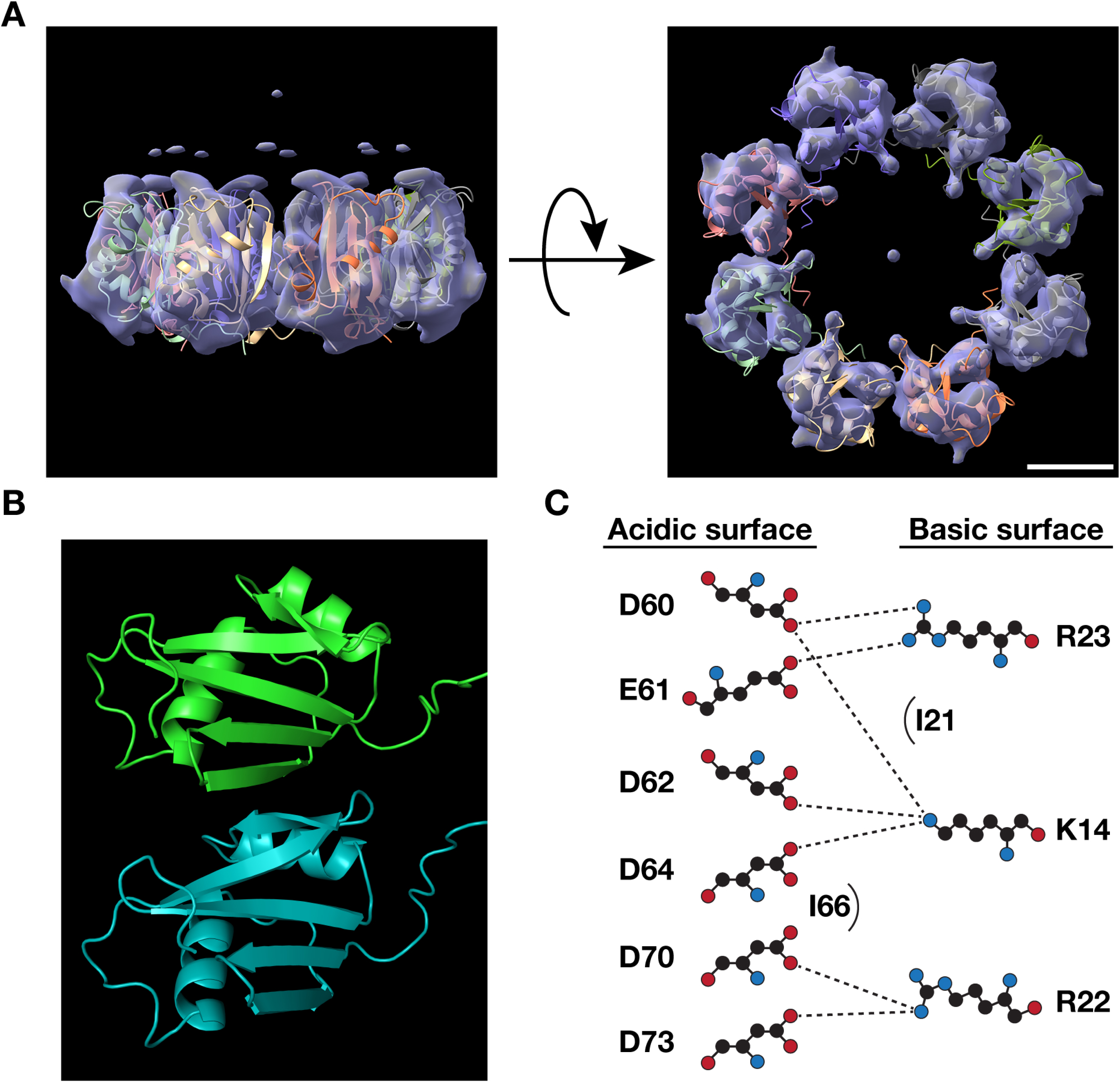
Structural model of the TFG ring complex based on cryo-EM. (A) Two views showing sharpened cryo-EM density maps of the TFG octameric ring complex shown as a transparent molecular surface, into which the TFG PB1 domain has been docked, leveraging Alphafold2 and ChimeraX. Bar, 5 nm. (B) Close-up view of the TFG PB1-PB1 domain interface, based on cryo-EM and X-ray crystallography. (C) Key salt bridge contacts and hydrophobic interactions at the predicted PB1-PB1 domain interface.

To validate the structural model, we conducted a series of mutagenesis studies, first incorporating individual amino acid changes into the TFG PB1 domain, which have been implicated previously in HSP (p.K14R, p.R22W, and p.I66T)^20,28,29^. Based on multi-angle light scattering and size exclusion chromatography studies, each variant reduced the molecular mass of the TFG PB1 domain, reflecting a defect in normal homo-oligomerization (**Figure 3A and 3B**). In addition, we separately neutralized a charged residue (D70) and mutated a non-polar residue (I21) within the PB1 domain, based on predictions that each plays an important role at the protomer interface by contributing to electrostatic and hydrophobic interactions, respectively. Similar to the impacts of the HSP-associated variants tested, both p.D70A and p.I21T disrupted the ability of the TFG PB1 domain to form octamers in solution (**Figure 3C and 3D**). In contrast, charge neutralization at K47, which is predicted to lie adjacent to the protomer interface, failed to impact oligomerization of the TFG PB1 domain (**Figure 3C and 3D**).

**Figure 3.**
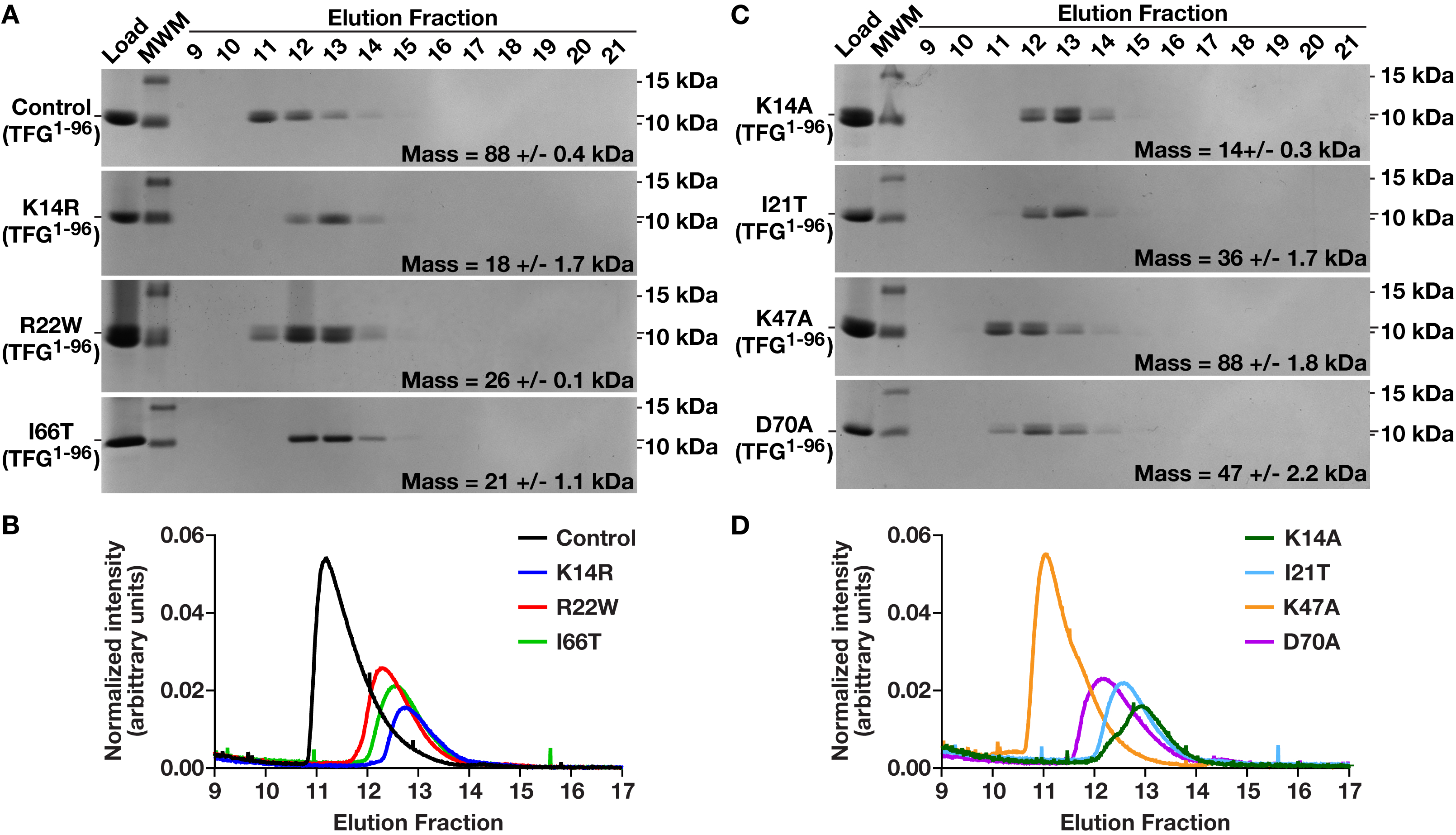
Validation of the PB1-PB1 domain interface in TFG ring complexes. (A and C) Recombinant isoforms of TFG (amino acids 1-96) were separated over a size exclusion chromatography column coupled to a multi-angle light scattering device. Eluted fractions were separated by SDS-PAGE and stained using Coomassie. The molar mass of each purified protein is shown (n=3 each). (B and D) Light scattering profiles are plotted for each protein, as indicated.

Notably, replacement of the lysyl side chain at K14 with a 3-carbon aliphatic straight-chain capped by a guanidino group, which does not affect its charge at physiological pH, was as disruptive as substitution with a methyl group lacking positive charge (**Figure 3C and 3D**). These data suggest that the TFG PB1 domain interface requires highly specific spacing to remain stable. However, in the context of the full length protein, none of the individual point mutations tested were sufficient to block oligomerization, likely due to contributions made by the coiled coil domain (**Figure S2A and S2B**). Consistent with this idea, we found that simultaneous mutations made in both the PB1 and coiled coil domains (p.K14A and p.R106C) impaired TFG oligomerization (**Figure 4A and 4B**). Taken together, our findings demonstrate that multiple HSP-associated mutations in the TFG PB1 domain weaken the same homotypic protomer interface to destabilize octameric ring assembly, thereby impairing TFG function.

**Figure 4.**
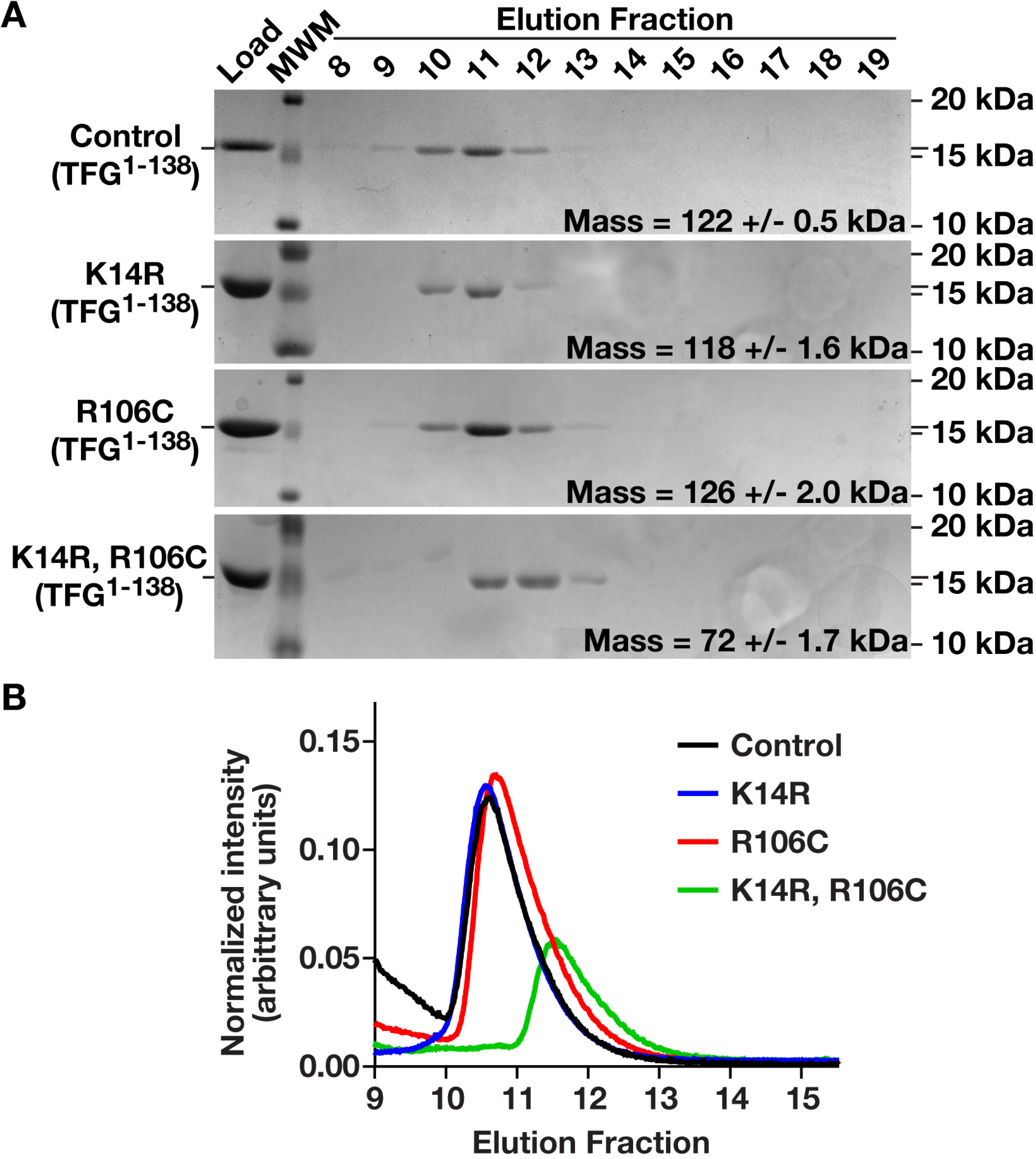
Both the PB1 domain and coiled coil motif of TFG contribute to oligomerization. (A) Recombinant isoforms of TFG (amino acids 1-138) were separated over a size exclusion chromatography column coupled to a multi-angle light scattering device. Eluted fractions were separated by SDS-PAGE and stained using Coomassie. The molar mass of each purified protein is shown (n=3 each). (B) Light scattering profiles are plotted for each protein, as indicated.

### A mutation in the TFG PB1 domain selectively impairs TFG stability in neurons

Pathological variants in TFG have been shown to alter its distribution in cells, although effects are heavily influenced by the specific domain impacted^9,16^. While the p.R106C mutation in the TFG coiled coil domain has been suggested to inhibit phase separation, resulting in a more diffuse distribution in cells^16,19^, the p.P285L mutation in the TFG disordered region leads to aggregate formation^9,21^. To determine how a mutation in the TFG PB1 domain affects localization, we focused on the p.R22W variant and examined dermal fibroblasts obtained from HSP patients expressing it in a homozygous manner^20^. Surprisingly, as compared to control fibroblasts, we found that the p.R22W mutation failed to impact TFG distribution (**Figure 5A and 5B**), its association with components of the COPII machinery (**Figure 5A**), or its expression level (**Figure 5C**), based on immunofluorescence and immunoblotting studies.

**Figure 5.**
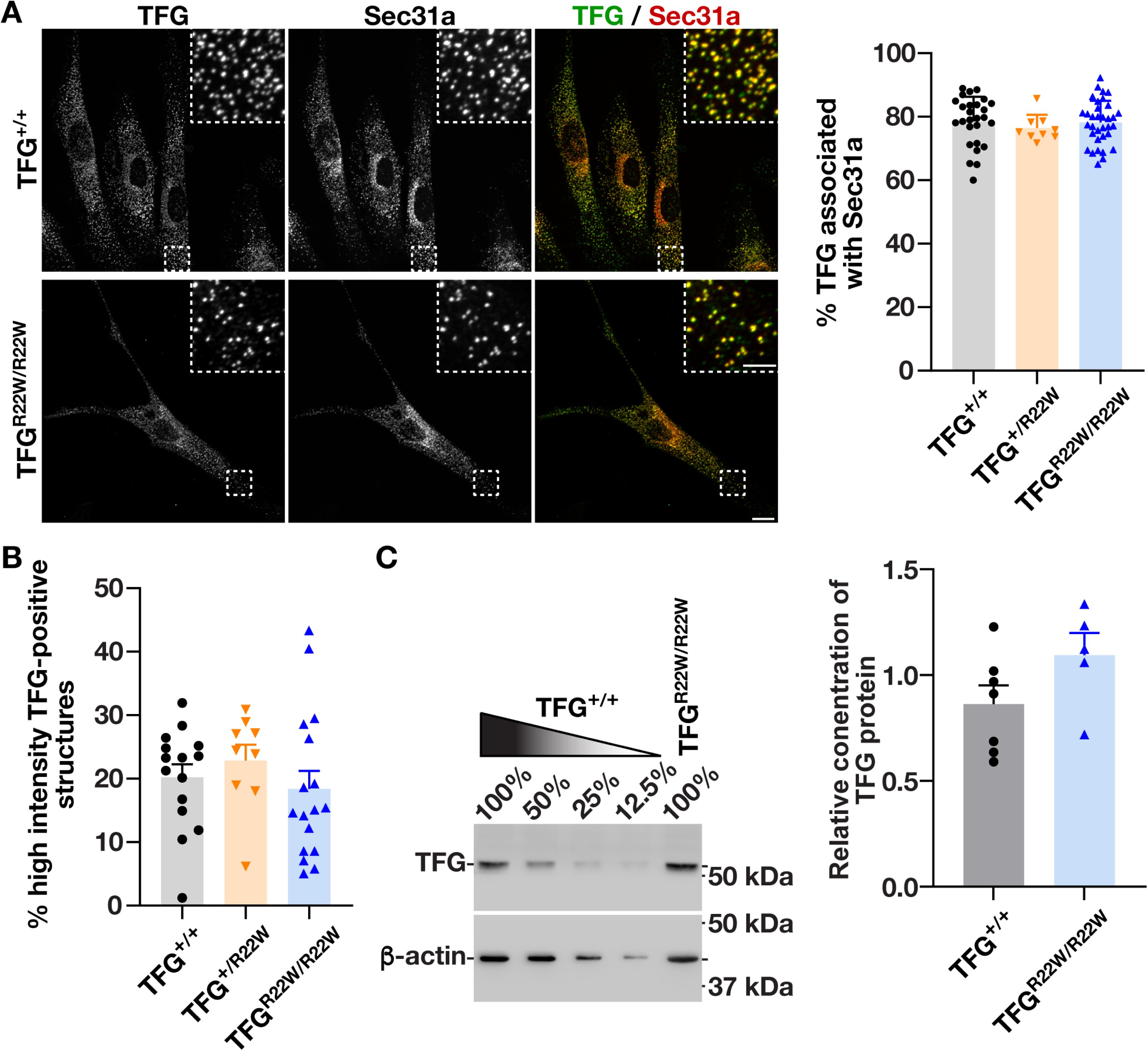
The p.R22W mutation in TFG fails to alter its expression or distribution in human patient-derived fibroblasts. (A) Control or homozygous mutant (TFG^R22W/R22W^) patient-derived fibroblasts were fixed and stained using antibodies directed against TFG (green) and Sec31A (red) and imaged using confocal microscopy. Maximum intensity projections of representative images are shown (left; n=10 cells each with 3 biological replicates each), with quantification of the percentage of TFG associated with Sec31A also indicated (right). Error bars represent mean +/- SEM, with no statistically significant differences found based on a one-way ANOVA and Tukey post hoc test. Scale bars, 10 μm and 5 μm (zoomed insets). (B) Quantification of the proportion of high-intensity TFG structures found in control and mutant patient-derived fibroblasts (n=10 cells each with 3 biological replicates). Error bars represent mean +/- SEM, with no statistically significant differences found based on a one-way ANOVA and Tukey post hoc test. (C) Representative immunoblots of extracts generated from control and mutant patient-derived fibroblasts, using antibodies directed against TFG and β-actin (left), with quantification of relative protein expression in control and mutant samples (right; n=7 each). Error bars represent mean +/- SEM, with no statistically significant differences found based on an independent two-sample t test.

Patients harboring the TFG p.R22W variant exhibit mainly neurological deficits, including thinning of the corpus callosum and axonal degeneration, raising the possibility that the effects of the mutation are restricted to cells of the CNS^20^. To investigate this idea, we first used CRISPR-Cas9 mediated genome editing to incorporate the p.R22W mutation into the germline of Sprague-Dawley rats. Kinematic gait analysis indicated significant and progressive motor dysfunction in homozygous TFG^R22W/R22W^ animals, highlighted by an increase in hindbody motion relative to littermate controls (**Figure S3A and S3B and Movie S1**). Additionally, as compared to control animals, homozygous mutant animals exhibited a hunched posture, showed an inability to sustain their weight, were unable to maintain tail tip height with age, and rapidly fell from an elevated revolving rod, all signs of progressive loss of muscle control (**Figure S3B-S3D**). These data suggest that TFG p.R22W animals represent a useful model for studying the causes and consequences of HSP.

To determine whether the p.R22W mutation alters the subcellular distribution of TFG or its expression levels within CNS tissue, we harvested the cerebral cortex from control and mutant animals and conducted a series of quantitative immunofluorescence studies following dissociation. Our findings indicated that TFG p.R22W continued to exhibit a punctate distribution, similar to wild-type TFG, but its intensity was significantly reduced in a subset of cells (**Figure 6A**). To determine the identity of this subpopulation, we co-stained cultures with antibodies directed against MAP2, a somatodendritic marker in multipolar neurons^30^. Strikingly, we found that reduced TFG p.R22W intensity was specifically associated with neuronal cells (**Figure 6A-6C**). Furthermore, based on immunoblotting of extracts generated from primary neurons grown in culture, we determined that TFG p.R22W levels were significantly reduced as compared to wild-type TFG, both in cell bodies and in neurites (**Figure 7A and Figure S4A-S4C**). Importantly, based on quantitative PCR studies, TFG transcript levels were unchanged relative to controls, strongly suggesting that the stability of TFG is adversely affected by the p.R22W mutation in neurons (**Figure S4D**). Consistent with our findings using patient-derived fibroblasts, primary rat embryonic fibroblasts from animals homozygous for the TFG p.R22W variant exhibited no differences in TFG localization or stability as compared to control fibroblasts, confirming that the impact of the mutation is neuron-specific (**Figure S5**).

**Figure 6.**
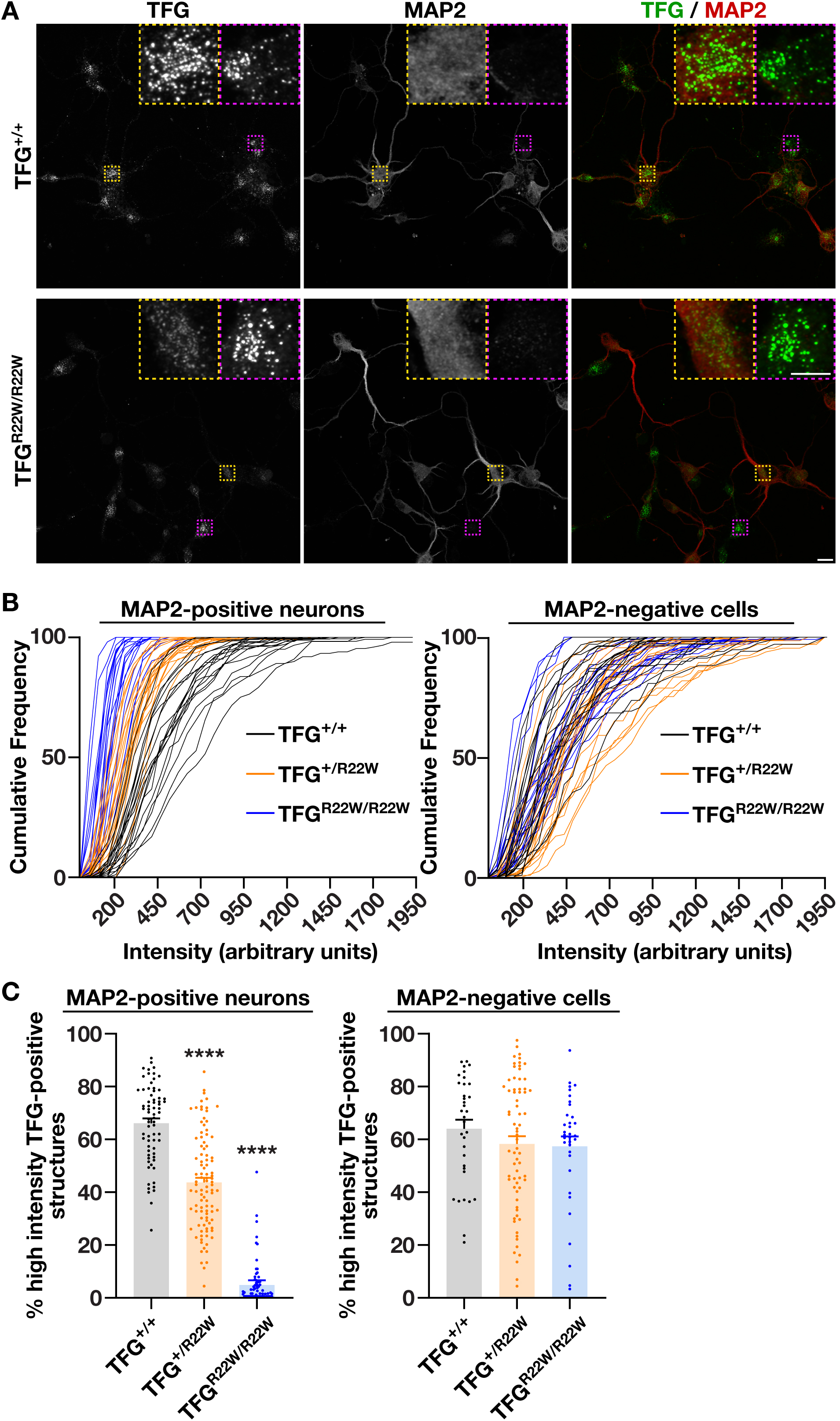
The p.R22W mutation in TFG impairs its expression specifically in MAP2-positive neurons. (A) Representative images of dissociated cortical tissue from control animals and animals homozygous for the TFG p.R22W mutation, which were grown in vitro for 2 days followed by fixation and staining using antibodies directed against TFG (green) and MAP2 (red). Maximum intensity projections are shown, with MAP2-positive cells highlighted by a yellow box, and MAP2-negative cells highlighted by a purple box. Scale bars, 10 μm and 5 μm (zoomed insets). (B) Relative cumulative histograms of the distribution of TFG intensities found in control and mutant MAP2-positive neurons (left) and MAP2-negative cells (right). (C) Quantification of the proportion of high-intensity TFG structures found in control and mutant MAP2-positive (left) and MAP2-negative (right) cells, following dissociation of cortices (n=30 cells each; 3 biological replicates each). Error bars represent mean +/- SEM. ****p < 0.0001, as calculated based on a one-way ANOVA and Tukey post hoc test.

**Figure 7.**
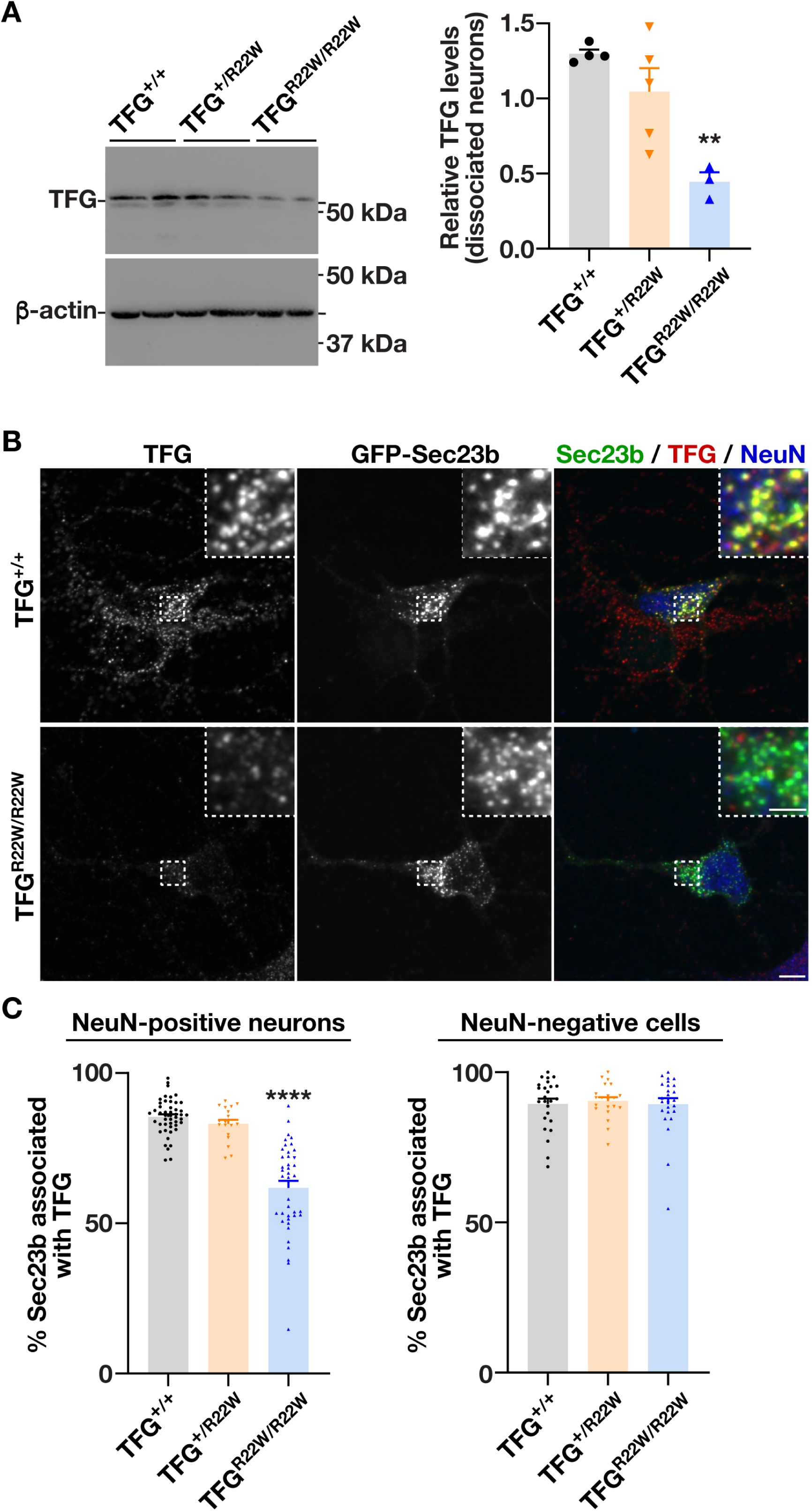
The p.R22W mutation in TFG impairs its distribution specifically in MAP2-positive neurons. (A) Representative immunoblots of extracts generated from control and mutant TFG p.R22W neurons grown in vitro for 14 days, using antibodies directed against TFG (top; left) and β-actin (bottom; left), with quantification of relative protein levels in control and mutant samples also shown (right; n=3 each). Error bars represent mean +/- SEM. **p < 0.01, as calculated based on a one-way ANOVA and Tukey post hoc test. (B) Representative images of dissociated cortical tissue from control animals and animals homozygous for the TFG p.R22W mutation, which were grown in vitro for 7 days following lentivirus transduction to express a GFP fusion to Sec23b (green), after which cells were fixed and stained using antibodies directed against TFG (red) and NeuN (blue). Maximum intensity projections are shown, with NeuN-positive neurons highlighted. Scale bars, 10 μm and 5 μm (zoomed insets). (C) Quantification of the percentage of GFP-Sec23b associated with TFG in NeuN-positive (left) and NeuN-negative (right) cells is indicated. Error bars represent mean +/- SEM. ****p < 0.0001, as calculated based on a one-way ANOVA and Tukey post hoc test.

To determine whether the reduced stability of TFG p.R22W impacts its ability to associate with components of the early secretory pathway, we transduced neurons with a virus expressing a GFP fusion to the COPII component Sec23b and examined their degree of overlap. These studies demonstrated that TFG p.R22W fails to accumulate normally at the ER-ERGIC interface, but only in neurons, as other cell types in primary cultures prepared from the cerebral cortex continued to show normal overlap with Sec23b (**Figure 7B and 7C**).

### The TFG p.R22W mutation slows cargo transport to the surface of axons

TFG has been shown to play essential roles in the trafficking of secretory and endocytic cargoes in multiple cell types^19,22,23,31^. To determine whether a mutation in its PB1 domain impairs cargo transport in neurons, we examined the cell surface accumulation of L1CAM, a cell adhesion molecule critical for neuronal migration and maintenance, in the presence and absence of the TFG p.R22W variant. These studies failed to demonstrate a significant impact of the mutation on the steady state distribution of L1CAM at the surface of axons (**Figure S6A and S6B**). Similarly, trafficking of L1CAM from the ER to the Golgi, as monitored using an inducible, synchronized release assay^16^, was not disrupted in the mutant neurons (**Figure 8A and 8B**). We therefore developed an approach to examine the kinetics of post-Golgi cargo trafficking, in which a HaloTag fusion to L1CAM was inducibly released from the ER and allowed to transit through the secretory pathway to the axonal plasma membrane over a period of 2.5 hours. Subsequently, cell impermeable and cell permeable HaloTag ligands, fused to distinct fluorescent dyes, were used in sequence to label external and internal populations of L1CAM respectively. Based on fluorescence intensity, we found that neurons expressing TFG p.R22W in a homozygous manner exhibited a kinetic delay in cell surface accumulation of L1CAM as compared to control neurons (**Figure 8C**). However, the defect in mutant neurons was relatively modest and failed to result in a reduction in axonal outgrowth as compared to controls (**Figure S6C**). These data suggest that a minimal threshold of functional TFG remains available in homozygous TFG p.R22W mutant neurons to support their activity. In contrast, further reduction in TFG levels by reducing gene dosage was not tolerated. Specifically, we found that compound heterozygous animals harboring one TFG p.R22W allele and one deletion allele (TFG^R22W/-^) could not be isolated, despite more than a dozen breeding attempts and more than 90 offspring genotyped. Together, our findings indicate that mutations in the PB1 domain impact TFG function distinctly from mutations associated with the coiled coil domain, despite both resulting in disruptions to TFG octameric ring assembly and clinical phenotypes associated with HSP.

**Figure 8.**
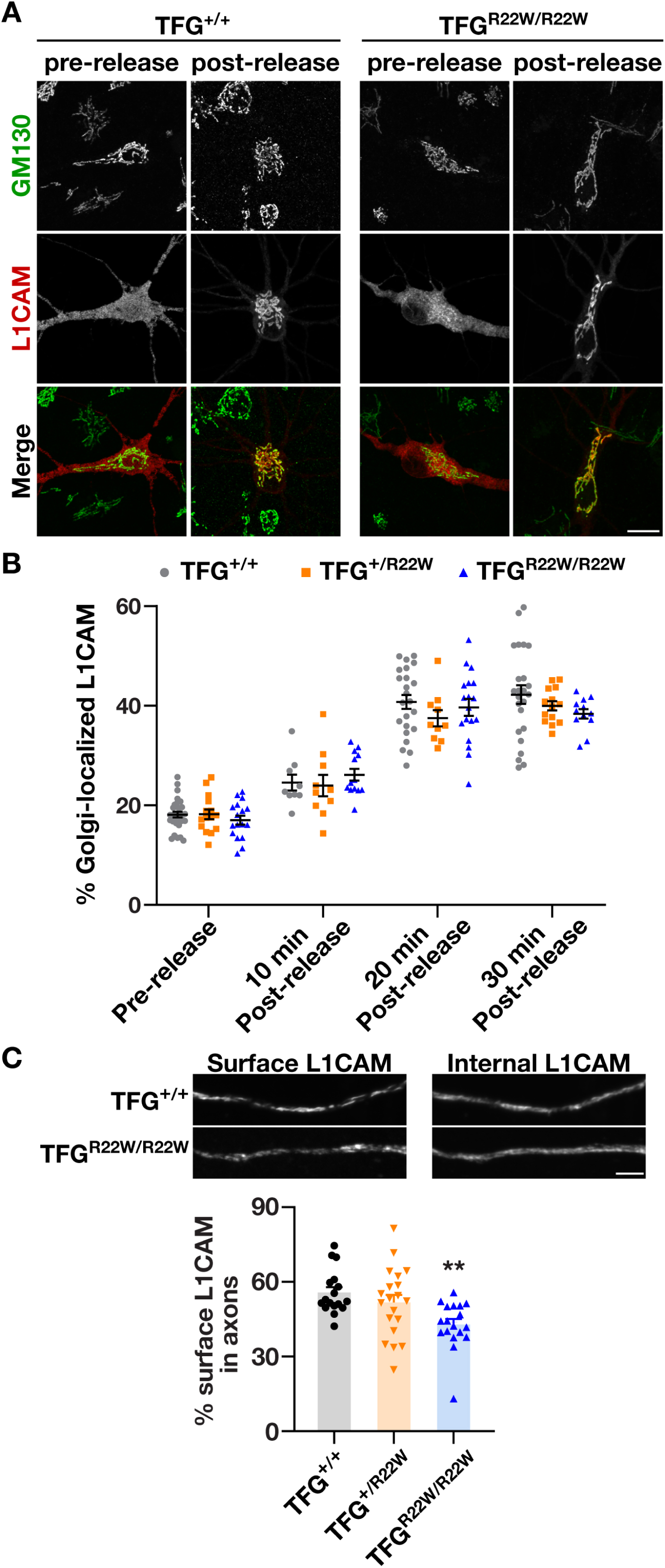
The p.R22W mutation in TFG impairs post-Golgi cargo trafficking in neurons. (A) Representative images of control and mutant TFG p.R22W primary rat cortical neurons transfected with a form of L1CAM that can be inducibly released from the ER, which has been fixed and stained using antibodies directed against GM130. Images were imaged using Spinning-disc confocal microscopy. Maximum intensity projections are shown before and 20 minutes after L1CAM release. Scale bar, 10 μm. (B) Quantification of the percentage of HaloTag-L1CAM associated with the Golgi over time following release from the ER in control neurons and neurons harboring the TFG p.R22W mutation (n=10 cells each per timepoint; 3 biological replicates each). Error bars represent mean +/- SEM. No statistically significant differences were identified, as calculated based on a one-way ANOVA and Tukey post hoc test. (C) Representative images of control and TFG p.R22W mutant primary cortical neurons transfected with an inducibly releasable form of HaloTag-L1CAM grown for 14 days in vitro and fixed 2.5 hours after the addition of DDS. Neurons were sequentially labeled with cell-impermeable, biotinylated HaloTag ligand (top; left), followed by a cell-permeable HaloTag ligand (top; right). Maximum intensity projections are shown. Quantification of the percentage of surface L1CAM as compared to total L1CAM intensity in axons from control and TFG p.R22W mutant neurons (n=20 cells each; 3 biological replicates each). Error bars represent mean +/- SEM. **p < 0.01, as calculated based on a one-way ANOVA and Tukey post hoc test. Scale bar, 5 μm.

## Discussion

PB1 domains have long been recognized for their ability to associate with one another, promoting the formation of macromolecular protein complexes, liquid condensates, and filamentous polymers that play diverse roles in cell physiology^32–35^. In animals, nine gene families encode PB1 domains, some of which heterodimerize exclusively with a singular cognate partner, while others exhibit extensive promiscuity^35^. Interestingly, the TFG PB1 domain has only been shown to interact with itself, predicting a highly distinctive subunit interface^36^. In general, specificity of PB1 domain interactions is directed by a unique network of electrostatic and hydrophobic interactions, generating high-affinity associations^35^. However, mechanistically defining the homotypic TFG PB1-PB1 interface has been challenging. In solution, single TFG particles exhibit a strongly preferred orientation along its eightfold axis, preventing three-dimensional structural determination^18^. Furthermore, while sedimentation enables recovery of approximately 90-nm TFG PB1 tubes, a high level of heterogeneity prevents a clear understanding of polymer formation^33^. Here, we combine X-ray crystallography, cryo-EM, and computational modeling to propose a high-probability model of TFG octameric ring complexes, highlighting the PB1-PB1 domain interface therein. Importantly, our findings demonstrate the mechanism by which several pathological mutations in the TFG PB1 domain lead to perturbations in TFG ring complex integrity, ultimately leading to neurodegeneration.

Despite the overall architecture of the TFG PB1 domain being reminiscent of others resolved previously^35^, the interface between subunits in TFG ring structures features a highly specific set of interactions, precluding co-assembly with alternative PB1 domains found in other proteins. Relatively subtle mutations, including the pathological p.K14R variant, are poorly tolerated, resulting in disease^28^. However, in this context, the presence of a coiled coil motif immediately downstream of the mutated PB1 domain promotes TFG oligomer assembly, albeit with an inability to form stable octameric rings. Similarly, a mutation in the TFG coiled coil domain also disrupts ring assembly^19^, leading to the idea that all mutations in the TFG amino-terminus predispose cortical neurons to degenerate in a similar manner. Our findings argue against such a model, showing instead that mutations in the PB1 domain and the coiled coil motif differentially impact TFG stability and its propensity to phase separate. Consistent with this idea, patients harboring variants in the PB1 domain (p.K14R, p.R22W, and p.I66T) exhibit distinct clinical presentations as compared to those expressing TFG p.R106C, with the latter causing more severe disease^8,20,28,29,37^. Remarkably, a comparison of our TFG p.R22W and p.R106C rodent models yields a similar conclusion, recapitulating key differences in motor dysfunction seen in human patients. Based on these data, we speculate that reduced stability of TFG caused by PB1 domain mutations is better tolerated by neurons as compared to a defect in condensate formation within the endocytic and early secretory pathways, which results from a mutation in the coiled coil motif, despite all amino-terminal mutations ultimately contributing to neuronal degeneration and a clinical spectrum of HSPs.

An alternative possibility is that the neuron-specific instability of TFG p.R22W restricts its overall physiological impact as compared to the effect of the p.R106C mutation, which is more broadly felt throughout the CNS. At a mechanistic level, it remains unclear how TFG p.R22W becomes selectively destabilized in neurons. In general, protein stability is dictated by a balance between protein synthesis and degradation rates, with brain tissue exhibiting longer protein lifetimes on average as compared to other organs^38^. Based on quantitative PCR studies, the TFG p.R22W variant is transcribed normally, suggesting that accelerated protein turnover likely accounts for its instability in neurons, potentially resulting from a defect in its ability to bind to a neuron-specific partner that normally shields TFG from ubiquitin-mediated degradation. There are several examples of factors that function in this protective manner, including 14-3-3, which binds to spastin and protects it from degradation, thereby facilitating neurite regeneration following injury^39^. Although it is unlikely that TFG binds to another protein via its homodimerization interface, mutations in this region lead to conformational changes, which may disrupt other interaction surfaces necessary for its stability. Future studies aimed at identifying additional TFG-binding proteins, whose expression is limited to neurons, will be important to validate this idea.

## Materials and Methods

### Recombinant protein expression and purification

All recombinant proteins were expressed as GST fusions using BL21(DE3) Escherichia coli (Sigma-Aldrich) and purified using glutathione-agarose beads, followed by removal of the GST tag. For size-exclusion chromatography, protein samples were either loaded onto a Wyatt WTC-030S5 or a Sepax SRT-C SEC-300 column, and eluted fractions were separated by Tricine-SDS page^40^. For multi-angle light scattering experiments, data were collected using a Wyatt miniDAWN TREOS three-angle light scattering detector, with UV absorbance and light scattering data collected every millisecond at a flow rate of 0.5 mL/min. Estimations of molecular mass were calculated using the ASTRA software^41^.

### Crystallization, structure determination and cryo-EM studies

The human TFG PB1 domain (residues 1-96; D60A/D62R) was concentrated to 6.6 g/L in 10 mM HEPES (pH 7.6), 100 mM NaCl, and 1 mM dithiothreitol, mixed in a 1:1 volumetric ratio with mother liquor (0.1 M Tris-HCl, pH 8.5, 8% PEG 8000) and crystallized by hanging drop vapor diffusion at 277 K. Cryoprotectant solution (0.1 M Tris-HCl, pH 8.5, 8% PEG 8000, 25% glycerol) was added directly to the crystals, and crystals were cryo-cooled in liquid nitrogen.

X-ray diffraction data collected at the Advanced Photon Source (Argonne National Labs) were indexed and scaled using HKL2000^42^. The structure of the protein was determined using molecular replacement leveraging a polyalanine model of the PB1 domain of Protein Kinase C (1WMH^43^) created using Chainsaw^44^ as the search model in Phaser^45^. The TFG PB1 domain model was refined through rounds of iterative manual building in Coot^46^ and refinement using REFMAC5^47^. Coordinate and structure factor files have been deposited in the Protein Data Bank (PDB Accession Code: 9E7C).

For cryo-EM studies, cryogenic grids were prepared using SPOTITON^48^. The system was loaded with 0.5 mg/mL TFG in 50 mM MES (pH 5.5) and 100 mM NaCl for blot-free freezing. Data were collected on a Titan Krios at 300 keV equipped with a Gatan K3 camera operated at 81,000X with a super-resolution pixel size of 0.557 Å/pixel. Movie frames were aligned with MotionCor2 with Fourier binning resulting in a final pixel size of 1.115 Å/pixel. Data were collected with a range of defocus from −2 to −4 μm. Two data sets were collected – one at 0° tilt with 1201 micrographs, and one at 30° tilt resulting in 1205 micrographs. Data were processed using cryoSPARC. Particles were picked using the template picker and good particles were selected from 2D class averages resulting in 144,305 particles from the tilted and untilted data combined. Ab-initio reconstruction in cryoSPARC was used to generate 3 initial maps, only one of which had sensible features. The 92,599 particles contributing to that map were refined against the ab initio map using non-uniform refinement with C8 symmetry to produce the final 6.76Å resolution map. The crystal structure of the TFG monomer was docked into all eight units of the map generated, leveraging AlphaFold2.

### Generation and characterization of CRISPR/Cas9-edited Sprague Dawley rats

Animal studies were approved by the Institutional Animal Care and Use Committee of the University of Wisconsin-Madison. Rodents were maintained in Innovive cages (12-hour light / 12-hour dark cycle) with free access to food and water, and health checks were performed daily. University and facility guidelines were adhered to for anesthesia and euthanasia.

CRISPR/Cas9-mediated genome editing of Sprague Dawley embryos was performed as described previously using a guide RNA (5’-CATTATGAATGGGAATTCGC-3’) and a single stranded a single-stranded oligodeoxynucleotide (5’-CTGGAGTCCACAATGAACGGACAGTTGGACCTAAGTGGGAAGCTGATTATCAAAGCTCAACTTGGAGAAGATATATGGCGAATTCCCATTCATAATGAAGACATTACTTATGATGAATTAGTGCTGATGATGCAGCGAGTATTCAGAGGA-3’) to incorporate a missense mutation encoding p.R22W. Edited animals were outcrossed with wild-type Sprague Dawley rats (Envigo).

Quantitative gait analysis was performed using a MotoRater apparatus (TSE systems) equipped with a flat platform, and analyzed using DeepLabCut and MATLAB as described previously^16^. For rotarod testing, rotation speed was set to 8 rpm and latency to fall was measured for each of five trials (one-minute break between trials).

### Growth of primary rat cortical neurons

Primary rat cortical neurons were collected as described previously^16^. Briefly, rat embryos (embryonic day 18.5) were decapitated, and the cortical brain tissue was isolated and incubated with Hibernate A supplemented with B-27 Plus. Tissue was subsequently digested with 0.25% Trypsin, followed by inactivation in Dulbecco’s modified Eagle’s medium (DMEM) supplemented with 10% fetal bovine serum. The digested cortices were mechanically dissociated by trituration in a micropipette tip and plated onto poly-D-lysine (PDL) coated glass coverslips. Neurons were cultured in Neurobasal A medium supplemented with B-27-Plus, penicillin, streptomycin, and GlutaMax at 37°C and 5% CO_2_. Transfection or transduction of cultured neurons was performed using Lipofectamine LTX or purified lentiviral particles.

### Confocal microscopy and immunofluorescence studies

For immunofluorescence studies, fibroblasts and primary rat cortical neurons were fixed using 4% paraformaldehyde and 4% glucose and permeabilized with 0.2%-0.5% TritonX-100 in PBS (except for the surface labeling experiments). Fixed cells were incubated with primary antibodies at 4°C overnight following blocking in 10% bovine serum albumin. After extensive washing, cells were incubated with secondary antibodies conjugated to fluorescent dyes for 1 hour at room temperature. Slides were again extensively washed and mounted into glass slides using VectaShield Antifade mounting medium for confocal imaging. Written informed consent was obtained from each participant who provided dermal fibroblasts, with the study approved by the Paris Necker Ethics Committee (France) and the Ethical Committee of the University of Khartoum Medical Campus (Sudan)^20^.

For trafficking assays, primary rat cortical neurons were transfected after four days in vitro (DIV4). At DIV14, neurons were first dye-labeled with JFX549-HaloTag ligand for 1 hour, and then washed 3 times with media. Dye-labeled neurons were then treated with dimer/dimer solubilizer (DDS) to release cargoes^16^. Neurons were fixed at various time points following treatment with DDS using 4% PFA and labeled as described above. For cell surface labeling of releasable HaloTag-L1CAM, neurons were only labeled after PFA fixation with JFX650-Biotin-HaloTag, followed by JFX549-Halotag ligand for 30 min at room temperature, with extensive washing after labeling with each.

For fluorescence imaging, a Nikon Eclipse Ti2-E spinning disc confocal microscope equipped with a Yokogawa CSU-W1 scan head, a 60X oil immersion objective lens, and an ORCA-Fusion BT sCMOS camera or an ImageXpress Micro 4 High-Content Imaging System (Molecular Devices) was used. Image analysis was conducted using Imaris or ImageJ.

### Statistical Methods

Reported p values were determined using GraphPad Prism. No data were excluded, and sample size was determined based on prior work.

## Data, Materials, and Software Availability

All study data are included in the article and/or supporting information.

## Acknowledgements

This work was supported in part by National Institute of Health Grants R35 GM134865 (to A.A.) and R01 NS124165 (to A.A.); the University of Wisconsin Carbone Cancer Center (UWCCC) Grant P30 CA014520; the Waisman Center Core Grant P50 HD105353. Support was also provided by the UWCCC Genome Editing and Animal Models Shared Resource, the UWCCC Flow Cytometry Laboratory, and the UW Optical Imaging Core facility. We thank Dr. Michael Hanna for assistance using AlphaFold2 and members of the Audhya lab for critically reading this manuscript.

## Declaration of Interests

The authors declare no conflicts of interest.

## Author Contributions

Z.Z., M.M.L., A.L.S., J.L.K., S.M.S., and A.A. designed research; Z.Z., M.M.L., A.L.S., B.B., P.R., T.N., I.K., A.R., J.R.A., and D.G., performed research; J.L.K., S.M.S., and A.A. contributed reagents/analytic tools; Z.Z., M.M.L., A.L.S., B.B., P.R., and A.A. analyzed data; and Z.Z., M.M.L., and A.A. wrote the paper.

## Notes

### Competing Interest Statement

The authors have declared no competing interest.

