## Supplemental Text for "Multiple roles for TFG ring complexes in neuronal cargo trafficking"

**Supplemental Figure Legends**

**Figure S1. Cryo-EM structure validation and metrics for the TFG single particle reconstruction.** (A) 2D class averages for the untilted and tilted datasets following cryo-EM imaging of the TFG amino-terminus. (B and C) Fourier shell correlation (FSC) curves are shown (B), as well as 3D FSC curves highlighting resolution anisotropy due to orientation bias in the 3D structure (C). Based on these data, sphericity = 0.639 out of 1; global resolution = 4.25 Å. (D) Euler angle plot suggests that the TFG amino-terminus adopts a strong preferred orientation (seen as lines representing the tilted and untilted data collection sessions).

**Figure S2. Individual point mutations within the TFG PB1 domain fail to impact its hydrodynamic profile.** (A) Recombinant isoforms of full-length TFG were separated over a size exclusion chromatography column coupled to a multi-angle light scattering device. Eluted fractions were separated by SDS-PAGE and stained using Coomassie (n=3 each). (B) Light scattering profiles are plotted for each protein, as indicated.

**Figure S3. Animals homozygous for the TFG p.R22W mutation exhibit mild gait deficits.** (A) Single frame from kinematic gait recording of a 25-week-old wild-type male animal (left) and a homozygous TFG p.R22W mutant male. Colored dots mark neural network-detected body parts. (B-D) Measurements of hindbody sway (B), body weight (C), and latency to falling from an elevated revolving rod (D) associated with 25-week wild type, heterozygous, and homozygous TFG p.R22W rats. Drawings above each graph schematize the quantified parameter. Triangles represent individual male animals; circles represent individual female animals. Error bars represent mean values +/- SEM. * $p<0.05$, ** $p<0.01$, **** $p<0.0001$, as calculated using one-way ANOVA with Tukey's post hoc test.

**Figure S4. The p.R22W mutation in TFG reduces its levels in neurites.** (A) Representative images of neurites associated with control and TFG p.R22W mutant neurons, which were grown in vitro for 2 days followed by fixation and staining using antibodies directed against TFG (green) and MAP2 (red). Maximum intensity projections are shown, with MAP2-positive cells highlighted by a yellow box, and MAP2-negative cells highlighted by a purple box. Scale bar, 10 µm. (B and C) Quantification of the proportion of high-intensity TFG structures (B) and their density (C) in control and mutant MAP2-positive neurons, following dissociation of cortices (n=30 cells each; 3 biological replicates each). Error bars represent mean +/- SEM. ****$p < 0.0001$, as calculated based on a one-way ANOVA and Tukey post hoc test. (D) Quantitative PCR was used to measure the expression of TFG in control and TFG p.R22W mutant neurons (n=3 biological replicates each). Error bars represent mean +/- SEM, with no statistically significant differences found based on a one-way ANOVA and Tukey post hoc test.

**Figure S5. The p.R22W mutation in TFG reduces its levels in neurites.** (A) Control or homozygous mutant TFG p.R22W rat embryonic fibroblasts were fixed and stained using antibodies directed against TFG (green) and GM130 (red) and imaged using confocal microscopy. Maximum intensity projections of representative images are shown (n=10 cells each; 3 biological replicates each). Scale bars, 10 µm and 5 µm (zoomed insets). (B) Relative cumulative histograms of the distribution of TFG intensities found in control and mutant rat embryonic fibroblasts. (C) Quantification of the proportion of high-intensity TFG structures found in control and mutant rat embryonic fibroblasts (n=10 cells each; 3 biological replicates each). Error bars represent mean +/- SEM, with no statistically significant differences found based on a one-way ANOVA and Tukey post hoc test. (D and E) Representative immunoblots of extracts generated from control and mutant rat embryonic fibroblasts, using antibodies directed against TFG and β-actin (D), with quantification of relative protein expression in control and mutant samples (E; n=4 each). Error bars represent mean +/- SEM, with no statistically significant differences found, based on an one-way ANOVA and Tukey post hoc test.

**Figure S6. Steady state levels of L1CAM on the axonal plasma membrane are unaffected by the TFG p.R22W mutation.** (A) Control and TFG p.R22W mutant primary rat neurons were grown in vitro for either 10 or 14 days, followed by fixation and staining with antibodies directed against L1CAM. Maximum intensity projections of representative images are shown (3 biological replicates each). Scale bar, 50 μm. (B) Quantification of surface L1CAM fluorescence intensity in control and TFG p.R22W mutant primary rat neurons at two different timepoints after plating. Error bars represent mean +/- SEM, with no statistically significant differences found, based on an one-way ANOVA and Tukey post hoc test. (C) Quantification of neurite length from control and TFG p.R22W mutant neurons grown in vitro for 2 or 4 days (n=19 cells each; 3 biological replicates each). Error bars represent mean +/- SEM, with no statistically significant differences found, based on an one-way ANOVA and Tukey post hoc test.

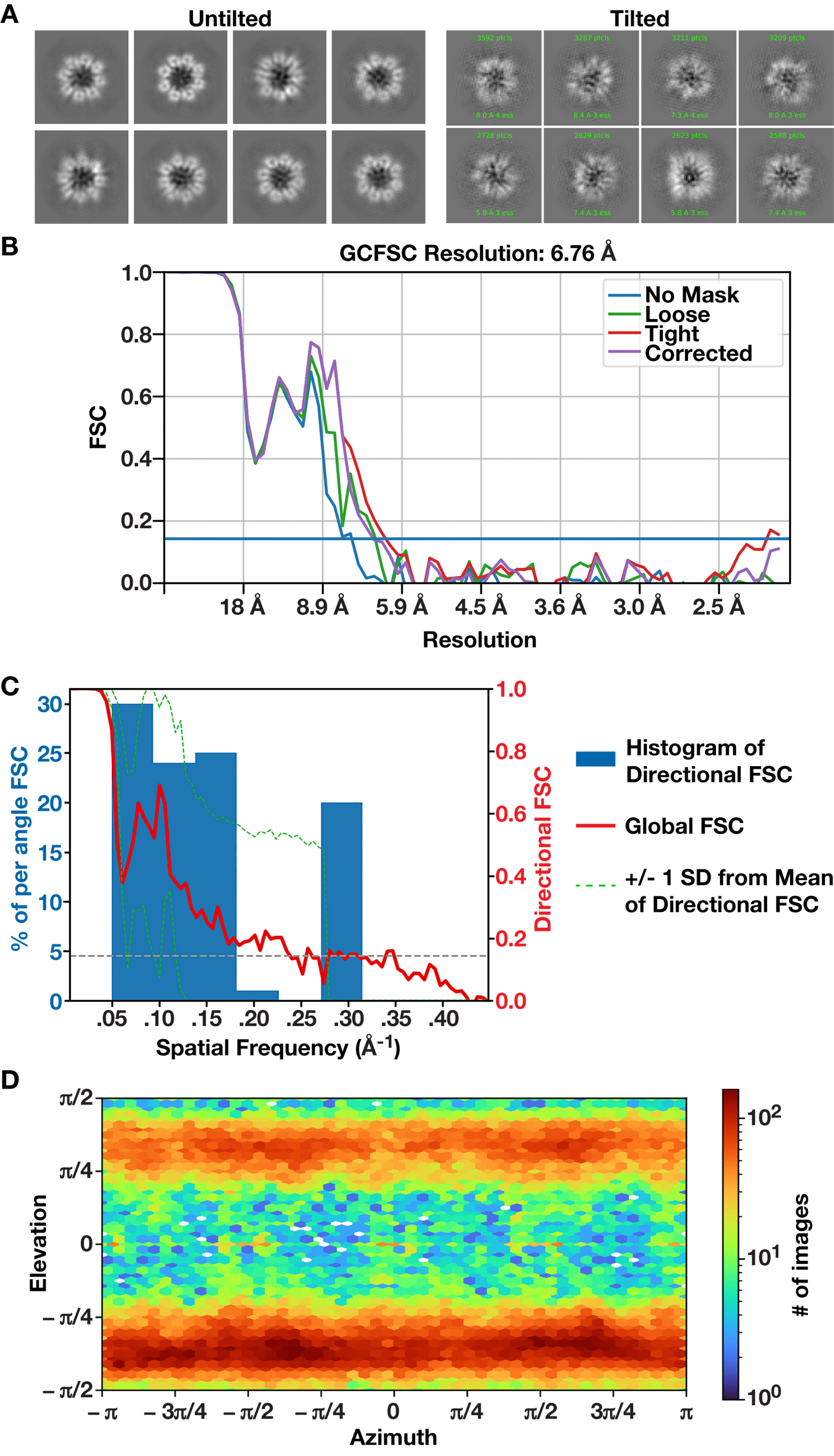
A
Untilted
Tilted
3592 ptcls
3287 ptcls
3211 ptcls
3209 ptcls
8.0 A 4 ess
8.4 A 3 ess
7.3 A 4 ess
8.0 A 3 ess
2728 ptcls
2629 ptcls
2623 ptcls
2588 ptcls
5.8 A 3 ess
7.4 A 3 ess
5.8 A 3 ess
7.4 A 3 ess
B
GCFSC Resolution: 6.76 Å
No Mask
Loose
Tight
Corrected
FSC
1.0
0.8
0.6
0.4
0.2
0.0
18 Å
8.9 Å
5.9 Å
4.5 Å
3.6 Å
3.0 Å
2.5 Å
Resolution
C
% of per angle FSC
30
25
20
15
10
5
0
1.0
0.8
0.6
0.4
0.2
0.0
Directional FSC
.05
.10
.15
.20
.25
.30
.35
.40
Spatial Frequency (Å$^{-1}$)
Histogram of Directional FSC
Global FSC
+/- 1 SD from Mean of Directional FSC
D
π/2
π/4
0
– π/4
– π/2
Elevation
– π
– 3π/4
– π/2
– π/4
0
π/4
π/2
3π/4
π
Azimuth
$10^2$
$10^1$
$10^0$
### of images

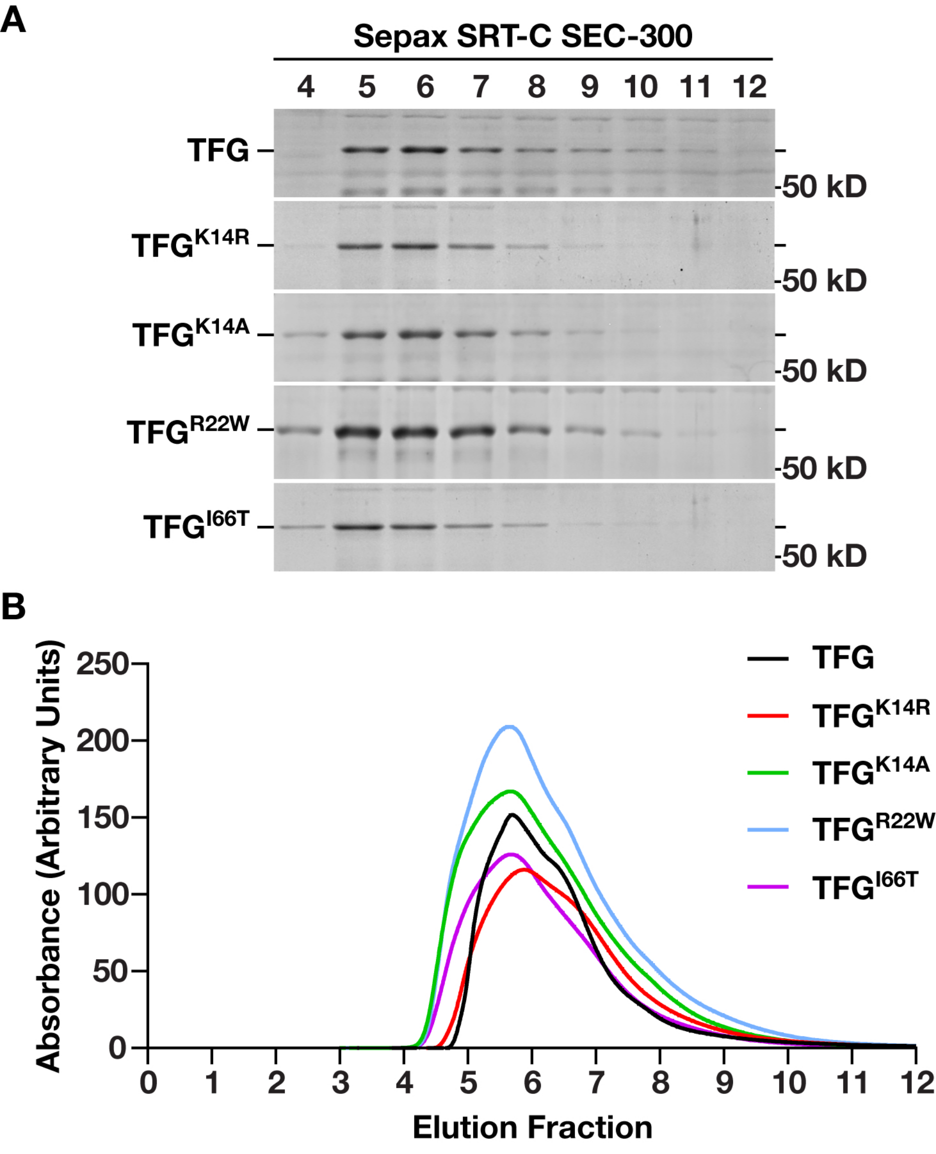
A
Sepax SRT-C SEC-300
4 5 6 7 8 9 10 11 12
TFG
TFGK14R
TFGK14A
TFGR22W
TFGI66T
-50 kD
B
Absorbance (Arbitrary Units)
250
200
150
100
50
0
0 1 2 3 4 5 6 7 8 9 10 11 12
Elution Fraction
TFG
TFGK14R
TFGK14A
TFGR22W
TFGI66T

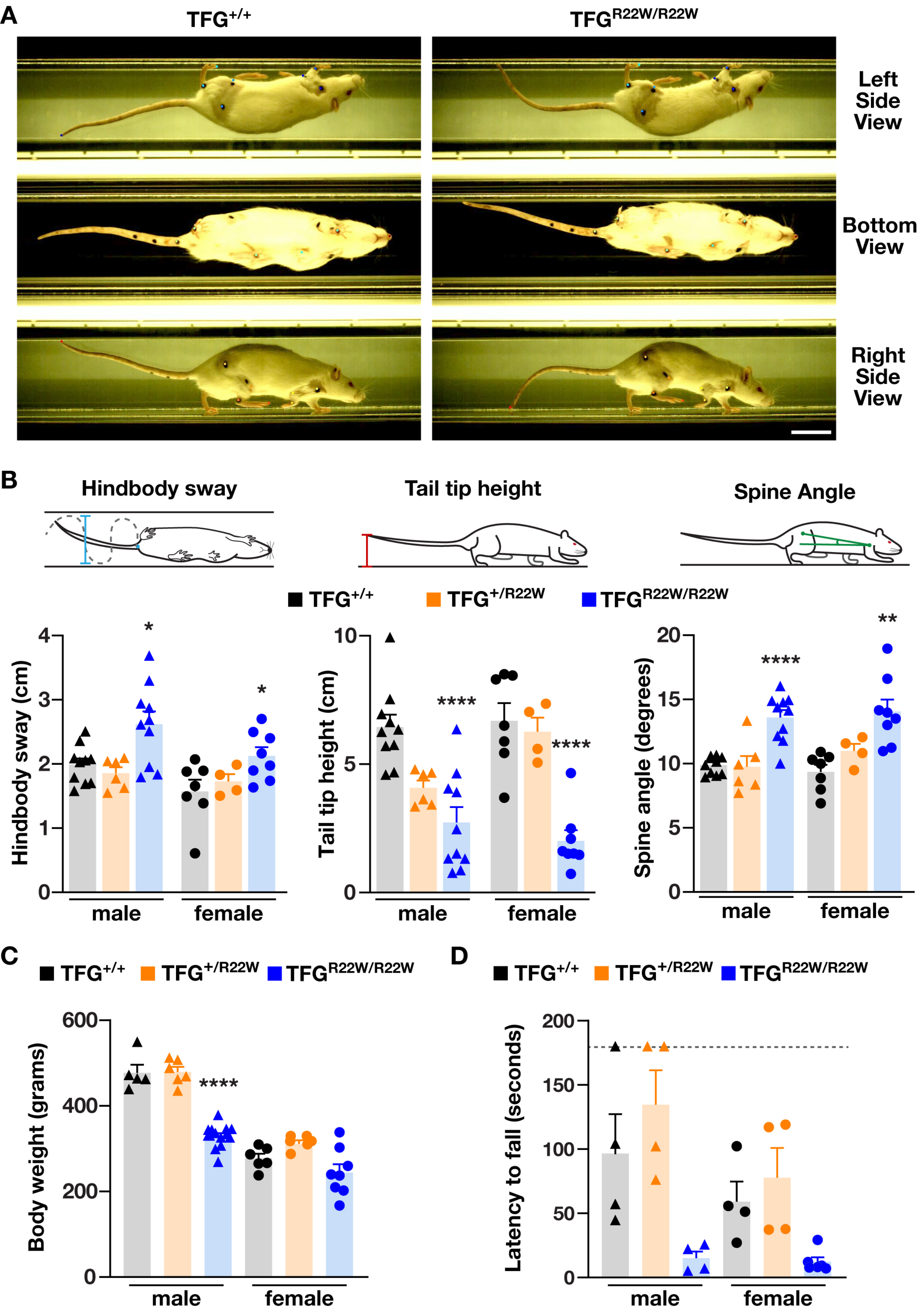
A
TFG+/+
TFGR22W/R22W
Left Side View
Bottom View
Right Side View
B
Hindbody sway
Tail tip height
Spine Angle
TFG+/+
TFG+/R22W
TFGR22W/R22W
Hindbody sway (cm)
Tail tip height (cm)
Spine angle (degrees)
male
female
C
TFG+/+
TFG+/R22W
TFGR22W/R22W
Body weight (grams)
male
female
D
TFG+/+
TFG+/R22W
TFGR22W/R22W
Latency to fall (seconds)
male
female

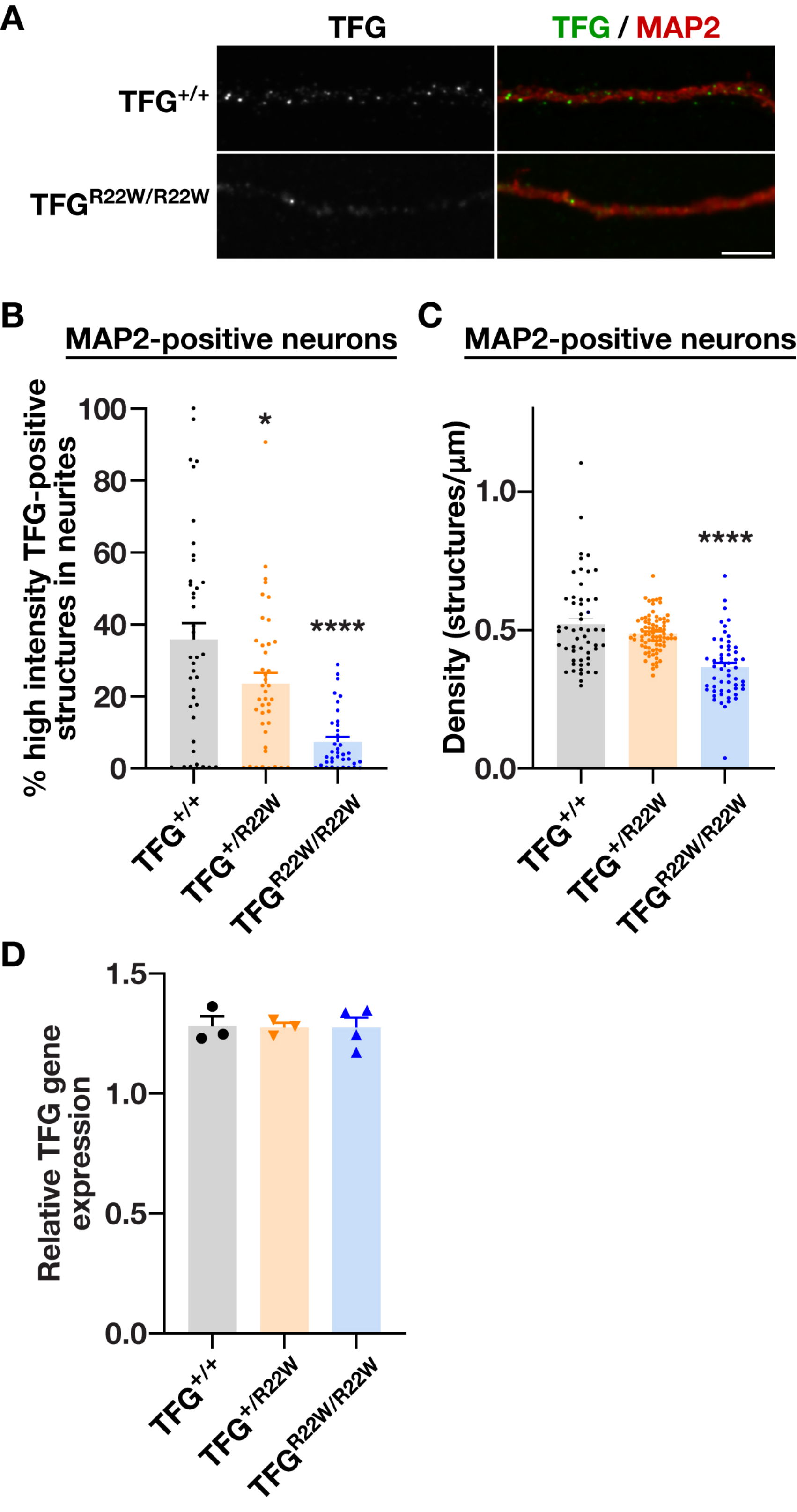
A
TFG
TFG / MAP2
TFG+/+
TFGR22W/R22W
B
MAP2-positive neurons
% high intensity TFG-positive structures in neurites
100
80
60
40
20
0
*
****
TFG+/+
TFG+/R22W
TFGR22W/R22W
C
MAP2-positive neurons
Density (structures/μm)
1.0
0.5
0.0
****
TFG+/+
TFG+/R22W
TFGR22W/R22W
D
Relative TFG gene expression
1.5
1.0
0.5
0.0
TFG+/+
TFG+/R22W
TFGR22W/R22W

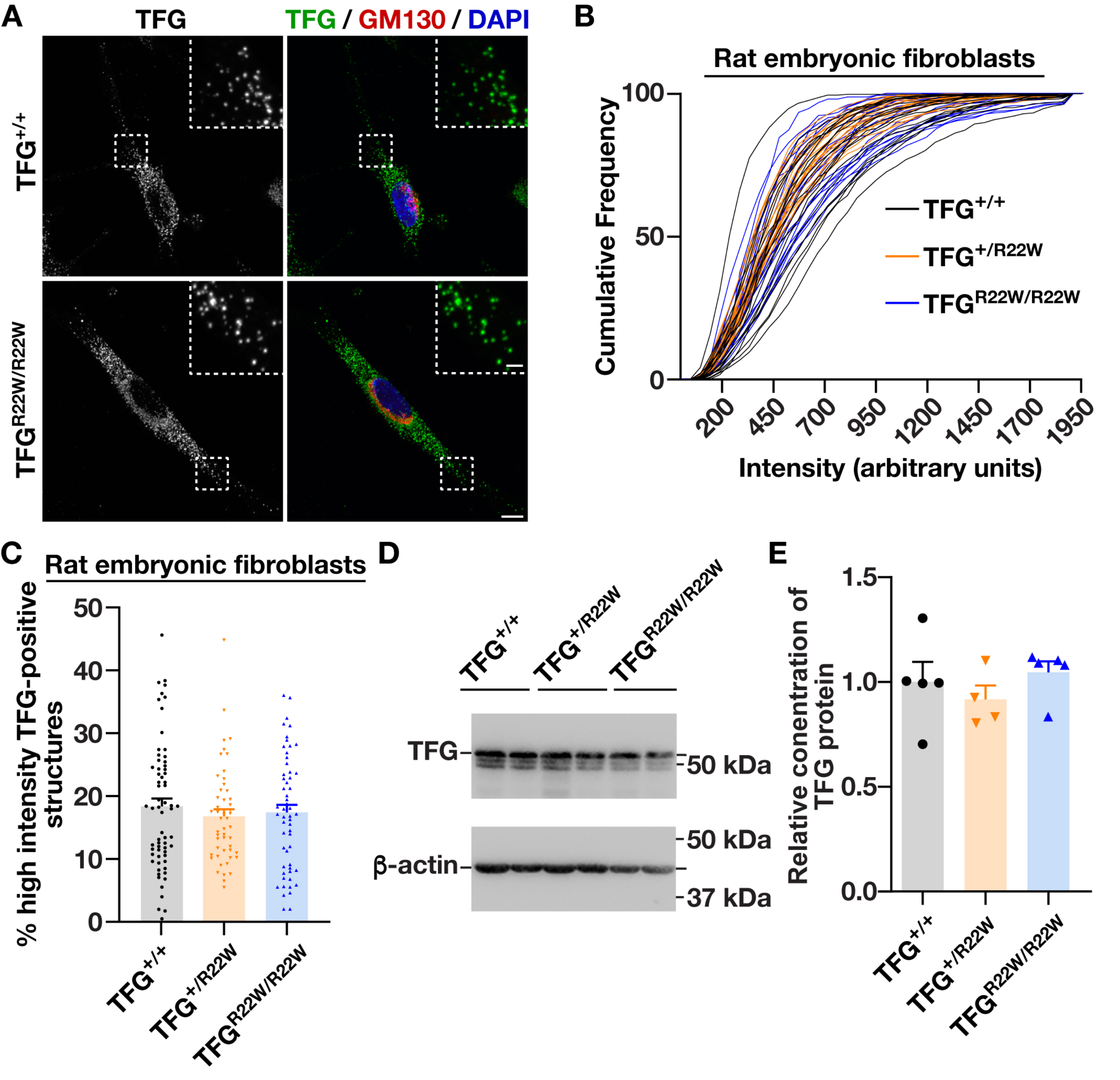

A
TFG
TFG / GM130 / DAPI
TFG+/+
TFGR22W/R22W
B
Rat embryonic fibroblasts
Cumulative Frequency
100
50
0
200
450
700
950
1200
1450
1700
1950
Intensity (arbitrary units)
TFG+/+
TFG+/R22W
TFGR22W/R22W
C
Rat embryonic fibroblasts
% high intensity TFG-positive structures
50
40
30
20
10
0
TFG+/+
TFG+/R22W
TFGR22W/R22W
D
TFG+/+
TFG+/R22W
TFGR22W/R22W
TFG
50 kDa
β-actin
50 kDa
37 kDa
E
Relative conentration of TFG protein
1.5
1.0
0.5
0.0
TFG+/+
TFG+/R22W
TFGR22W/R22W

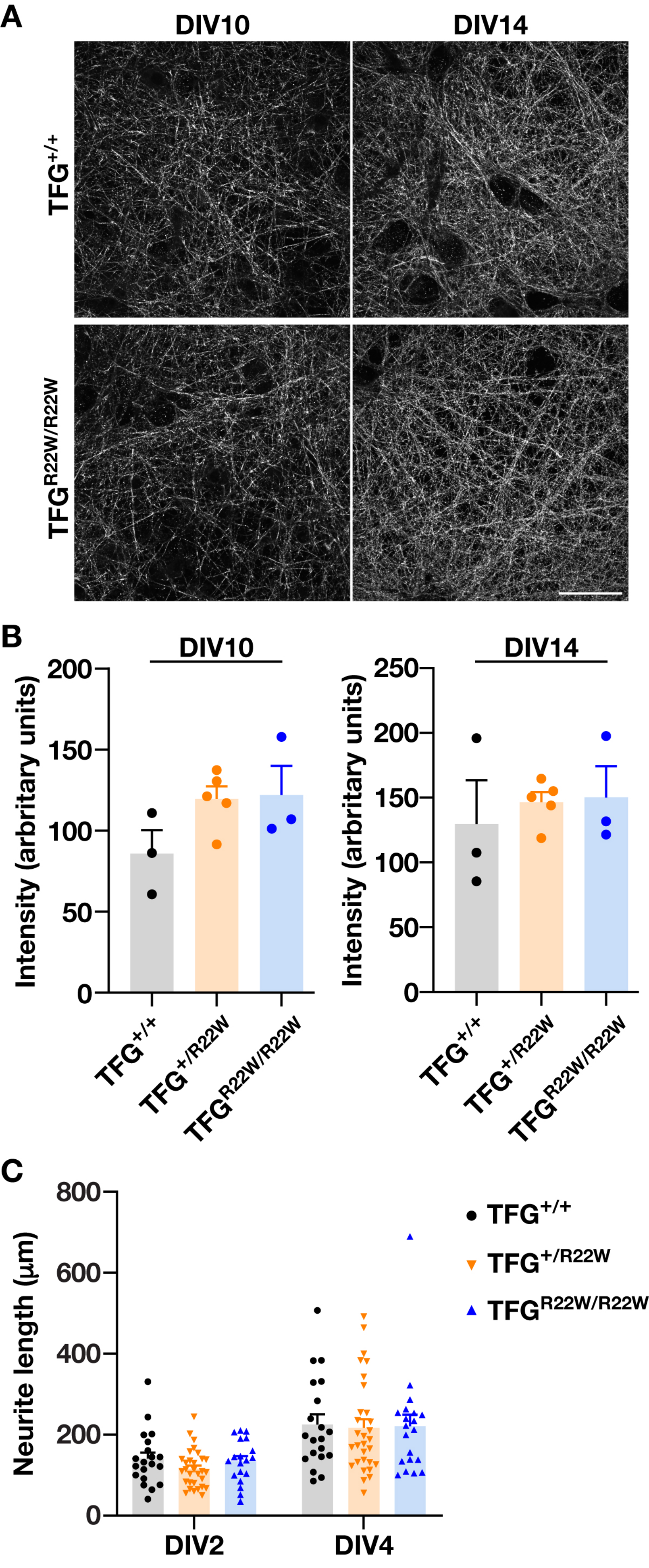
A
DIV10
DIV14
TFG+/+
TFGR22W/R22W
B
DIV10
DIV14
Intensity (arbitrary units)
200
150
100
50
0
250
TFG+/+
TFG+/R22W
TFGR22W/R22W
C
Neurite length (µm)
800
600
400
200
0
DIV2
DIV4
TFG+/+
TFG+/R22W
TFGR22W/R22W

**Supplemental Movie Legend**

**Movie S1. Representative videos of 25-week old male rats.** Multiple views of animals as they traverse a flat platform. Specific anatomical features of interest mapped by a neural network are represented as colored points. Control animals (TFG$^{+/+}$, left), animals with one mutant allele (TFG$^{R22W/+}$, middle), and homozygous TFG$^{R22W/R22W}$ animals (right) are shown.
